# Cancer-specific sialylation of insulin-like growth factor 1 receptor impairs therapeutic antibody binding and efficacy

**DOI:** 10.1101/2025.10.17.682592

**Authors:** Ayon Ahmed Hassan, Qiushi Chen, Katie Sze Wai Fung, Sally Kit Yan To, Lydia Wai Ting Cheung, Chenhui Zhang, Nan Liu, Chun Nan Lok, Chi-Ming Che, Philip Pun Ching Ip, Xuechen Li, Alice Sze Tsai Wong

**Author notes:** Corresponding authors: Alice S. T. Wong,; Xuechen Li. Equal contribution (Ayon Ahmed Hassan, Qiushi Chen).

## Abstract

Despite extensive efforts to develop insulin-like growth factor (IGF1R)-targeted therapies for various malignancies, none has received clinical approval in the past two decades. Here, we reveal that N-glycan sialylation significantly decreases recognition by the humanized monoclonal anti-IGF1R antibody ganitumab across various cancer types, reducing its efficacy both *in vitro* and *in vivo*. Sialoforms of IGF1R are virtually absent in normal cells, indicating that the modification is tumor-specific. Pharmacological inhibition of sialyltransferases significantly sensitizes metastatic tumors to ganitumab in a ganitumab-resistant ovarian cancer model. Enzymatic removal of sialic acids from tissue sections resulted in marked enhancement in antibody binding to ovarian cancer patient tumors, but not normal tissues. Upregulation of α2-6 sialyltransferase ST6GAL1 in tumor tissues was found to be responsible for sialylation of IGF1R. Consequently, ST6GAL1-high tumors were more likely to benefit from desialylation-mediated enhancement of ganitumab binding. Furthermore, through comprehensive glycoproteomics analysis, structural prediction, and molecular dynamics simulation, we identify Asn-607 (N607) as a crucial site harboring sialylated glycans. Mechanistically, N607 glycosylation destabilizes the IGF1R-ganitumab complex. Overexpression of IGF1R Asn-607-Gln (N607Q) mutant in IGF1R-knockout cancer cells increases ganitumab efficacy compared to wild-type IGF1R *in vivo*. Taken together, these findings highlight sialylation as a common barrier in IGF1R-targeted therapies and provide crucial insights for therapy enhancement in cancer and patient stratification for future clinical trials.

**Graphical Abstract:** 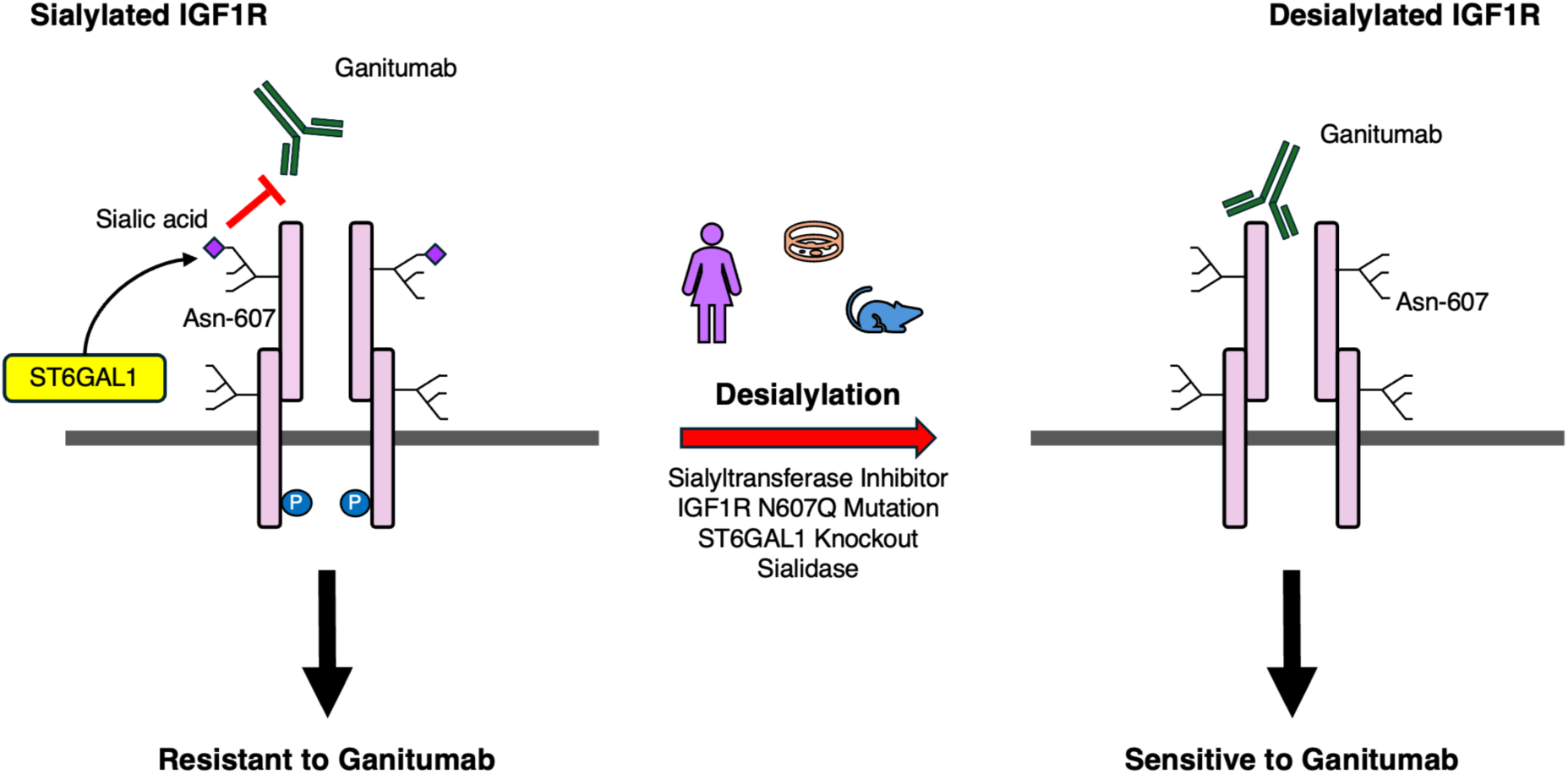

**Synopsis:** α2-6 sialylation of IGF1R Asn-607 by ST6GAL1 is prevalent in cancer. This modification disrupts the interaction of IGF1R with therapeutic mAb ganitumab. Removal of sialylation augments ganitumab efficacy.

## Introduction

The insulin-like growth factor 1 receptor (IGF1R) is a key receptor tyrosine kinase involved in essential processes such as growth, survival, and metastasis in cancer. Dysregulated IGF1R signaling has been observed in at least 15 different types of cancers, including breast, lung, colorectal, prostate, and ovarian cancers^1^. Over the past two decades, extensive efforts have been dedicated to developing therapeutic agents against this receptor, making it one of the most well-studied molecular targets in cancer research. Despite promising initial results of IGF1R inhibition in certain cancers, the translation to sustained success has been challenging^1–3^. From 2000 to 2021, over 16 IGF1R inhibitors/monoclonal antibodies (mAbs) were assessed in 183 clinical trials involving more than 12,000 cancer patients, with total costs exceeding $1.6 billion^2^. While anti-IGF1R mAbs, such as ganitumab and dalotuzumab, have reached Phase III clinical trials, all of them yielded sub-optimal outcomes^1^. Currently, ganitumab remains the only one still in Phase III trials^1^. Treatment insensitivity have commonly been attributed to compensatory pathways, such as PI3K, MAPK, and insulin receptor activation, but co-targeting these pathways have proven ineffective in enhancing therapeutic response^1–3^. These indicate additional layers of regulation not adequately targeted by current therapies, in particular the role of post-translational modifications. It is worth noting that no trials for ovarian cancer have advanced beyond Phase II, suggesting that this cancer type could be a valuable model for studying such intrinsic resistance to IGF1R-targeting therapies^4–6^. Ovarian cancer is particularly challenging to treat due to late-stage detection and high recurrence rates following standard therapies^7^. While FDA approved PARP inhibitors have shown promise in benefiting a subset of patients with homologous recombination deficiencies in recent years, ovarian cancer still remains the most lethal gynecological malignancy^8^. Therefore, there is an urgent need to explore alternative approaches to effectively address this disease. Our previous study has highlighted a significant role of glyco-modifications in the peritoneal dissemination of ovarian cancer^9^, a phenomenon that is commonly linked with therapy resistance^10^.

Aberrant glycosylation, an important hallmark of cancer, plays a crucial role in shaping protein dynamics and functions^11,12^. This post-translational modification involves addition of glycan moieties to proteins, which alters their chemical and biological properties by impacting flexibility, movement, and charges. Protein glycosylation was also recently shown to influence its interaction with antibodies. For example, N-glycans on the immune checkpoint PD-L1 can impede antibody binding, and their removal could enhance the accuracy of PD-LI histologic quantification, thereby improving predictive value for clinical outcomes^13^. Previous research has indicated that specific glycan modifications such as sialylation of receptor tyrosine kinases can affect the efficacy of targeted therapies, for example, gefitinib (targeting EGFR) in lung cancer^14^ and trastuzumab (targeting ERBB2) in gastric cancer^15^, but no studies have yet unveiled a precise mechanism that can explain clinical therapeutic resistance. Most studies to date lack one or more crucial information related to therapeutic inefficacy including (i) the precise site(s) of glycosylation, (ii) specific glycoforms involved, and (iii) their regulation. This poses a significant barrier to translating glycosylation-related discoveries into practical therapeutic strategies. Moreover, these studies are conducted primarily *in vitro* using cellular or biochemical assays and lack animal models and patient analysis.

We hypothesize that glycans on IGF1R may impede the efficacy of mAb-based IGF1R-targeted therapeutics, leading to therapeutic resistance. In this study, we make a notable advancement by providing a mechanistic explanation that directly links site-specific sialylation of IGF1R to resistance against ganitumab. The presence of sialylated IGF1R is observed in prevalent cancer types that have faced challenges in ganitumab trials, suggesting a widespread implication of this modification in diverse cancers. We identified a tumor-specific sialoform of IGF1R and demonstrated that ST6GAL1-mediated N-glycan sialylation on N607 is responsible for ganitumab resistance. Combined with molecular dynamics simulation, these provide a mechanistic explanation for therapy failure in the clinic. Together, our findings suggest that targeting the sialylation of IGF1R could provide a novel approach to enhance the efficacy of IGF1R-targeted therapies, and provide a rationale for stratifying patients that are likely to benefit from this treatment.

## Results

### Desialylation sensitizes ovarian cancer cells to anti-IGF1R mAb ganitumab

Sialylation, the addition of sialic acid to terminal glycan chains, is a critical glycosylation modification on cell surface proteins that is upregulated in cancers^16^. To begin, we first assessed the impact of sialylation on ganitumab treatment in a xenograft model of metastatic ovarian cancer. Mice were intraperitoneally injected with SKOV-3 cells and subsequently treated with ganitumab alone or in combination with a chemical sialyltransferase inhibitor (STI), 3Fax-Peracetyl Neu5Ac. Ganitumab monotherapy did not significantly affect metastasis compared to IgG-treated control group, indicating that our model is able to recapitulate the resistance observed in patients. On the other hand, mice receiving the combined ganitumab and STI treatment exhibited a marked reduction in tumor growth, as evidenced by significantly decreased signal intensity in bioluminescence imaging and a reduced number of tumor nodules on the mesenteries **(Figure 1a)**. To investigate the mechanism, we first tested if sialylation can directly influence the interaction of ganitumab with IGF1R by immunoprecipitating sialidase-treated IGF1R from ovarian cancer cell lysates using ganitumab. Treatment with sialidase significantly increased the amount of IGF1R in the IP eluate, suggesting that desialylation could specifically promote IGF1R’s physical binding to ganitumab **(Figure 1b)**. We also tested the effect of another terminal glyco-modification, fucosylation, on ganitumab-IGF1R interaction. No significant difference was observed compared to control **(Supplementary Figure S1a)**, suggesting that the effect is specific to sialic acid. There are reports that loss of glycosylation alters the cell surface availability of proteins and hybridization of IGF1R with insulin receptor (INSR). However, we observed that desialylation has no effect on IGF1R cell surface localization, activation, or hybridization with INSR **(Supplementary Figures S2a-c)**. Taken together, these suggest that the results observed in our xenograft model is likely due to direct effect of sialic acid on IGF1R-ganitumab complex.

**Figure 1.**
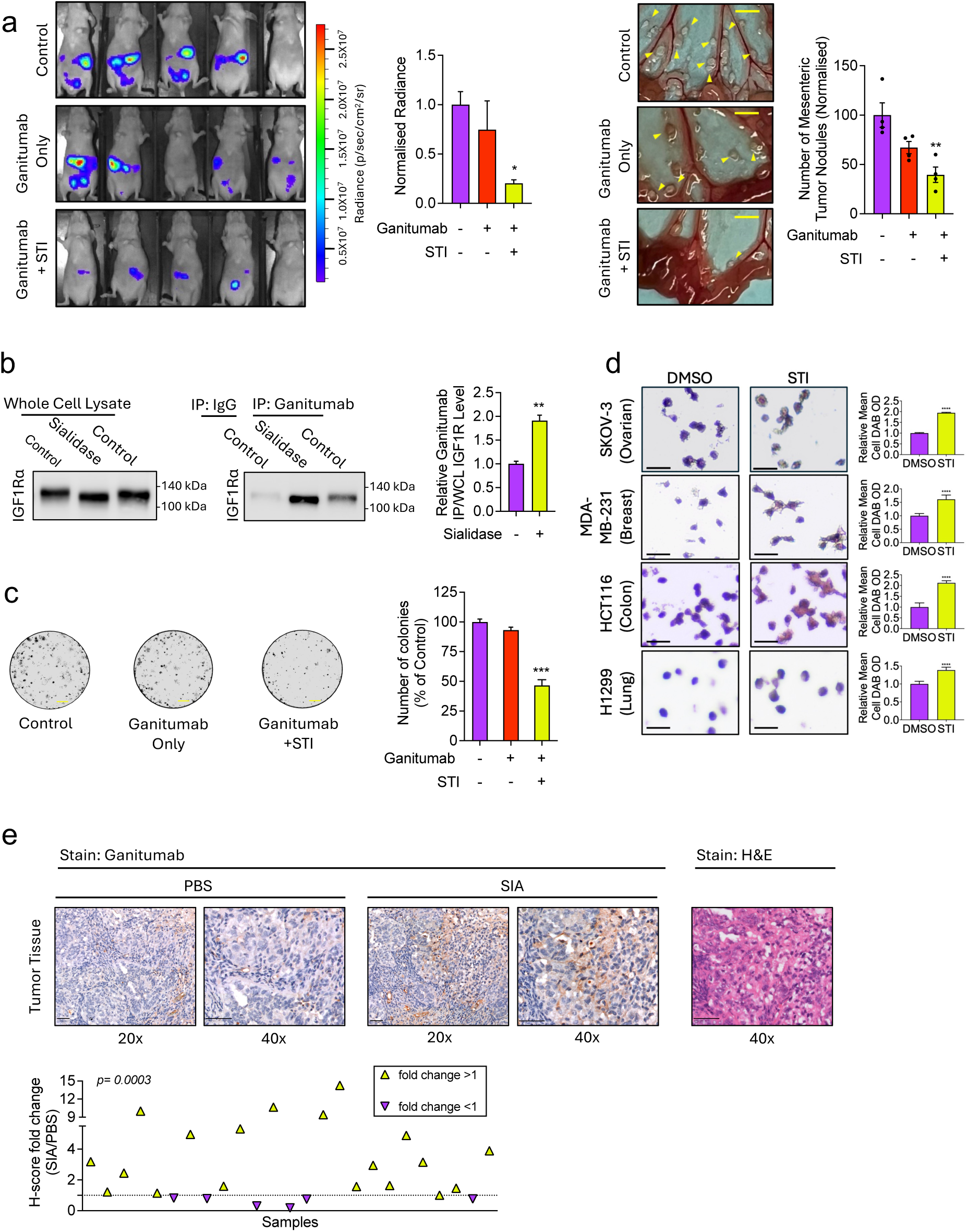
Desialylation sensitizes ovarian cancer cells to anti-IGF1R mAb ganitumab. **(a)** Bioluminescence imaging and mesenteric tumor nodules in a xenograft model of ovarian cancer cells treated with ganitumab only or in combination with sialyltransferase inhibitor (STI); n= 5 per group; scale bar= 0.25cm. **(b)** Immunoprecipitation of sialidase-treated insulin-like growth factor 1 receptor (IGF1R) with ganitumab. **(c)** Clonogenic cell survival assay of ovarian cancer cells treated with ganitumab only or in combination with (STI); scale bar= 0.5mm. **(d)** Average ganitumab signal intensity per cell (mean cell DAB OD) in STI-treated ovarian cancer, breast cancer, colon cancer, and lung cancer cell blocks by IHC; scale bar= 25μm. **(e)** Ganitumab H-score of ovarian cancer (n=25) FFPE tissues after sialidase (SIA) digestion; representative image of the staining and H&E staining of the tissues shown at 20x and 40x magnification. Scale bar = 50μm.

Next, we investigated whether these glycosylation modifications affect the sensitivity of ovarian cancer cells to ganitumab in a clonogenic cell survival assay. Ganitumab alone did not have significant effect on colony formation of SKOV-3 cells, indicating their inherent resistance to ganitumab **(Figure 1c).** However, the presence of STI significantly decreased the ability of ovarian cancer cells to form colonies under ganitumab treatment. No notable difference was observed with 2F-Peracetyl-Fucose **(Supplementary Figure S1b)**, a fucosyltransferase inhibitor, suggesting that sialylation, but not fucosylation, could reduce ganitumab sensitivity. Consistently, treatment with STI increased the immunohistochemical (IHC) staining intensity of ganitumab in ovarian cancer cells **(Figure 1d)**, further supporting the notion that sialylation impedes ganitumab’s recognition of IGF1R. We also tested the effect of sialylation on ganitumab-IGF1R interaction in aggressive cell lines of other common cancer types including breast (MDA-MB-231), colon, (HCT116), and lung (H1299) cancers. Consistent with our findings in ovarian cancer cells, IHC staining demonstrated that STI treatment significantly increased ganitumab binding in these cell lines **(Figure 1d)**. There was no significant difference for STI-treated normal human ovarian surface epithelial cells **(Supplementary Figure S1c)**. STI treatment also increased binding of another anti-IGF1R therapeutic antibody, dalotuzumab, to ovarian cancer cells **(Supplementary Figure S1d)**. Sialylation reduction by STI treatment was confirmed by staining with SNA lectin, which binds to α2-6-linked sialic acid and weakly to α2-3-linked sialic acids **(Supplementary Figure S1e)**. Together, these results suggest that sialylation of IGF1R may be prevalent across various cancer types, and removal of it could enhance recognition of IGF1R by therapeutic antibodies.

To assess the clinical relevance of our findings, we obtained formalin-fixed paraffin-embedded (FFPE) sections from ovarian cancer patients and subjected them to sialidase digestion followed by immunostaining with ganitumab. A significant enhancement in ganitumab signal intensity was observed following desialylation in tumor samples **(Figure 1e and Supplementary Figure S1g)**, while no discernible difference was noted in normal ovarian tissues **(Supplementary Figures S1e and S1h)**, providing direct evidence that tumor-specific sialylation of IGF1R could impede IGF1R-ganitumab interaction.

### IGF1R is modified with α2-6-linked sialylated N-glycans

In humans, sialic acids are observed as terminal structures on N- or O-glycans, attached via α2-3, α2-6, or α2-8 linkages^17^. The type of glycan sialic acids decorate and the type of linkage determines the biological behavior of the molecule^18^. To elucidate the structure of sialylated glycans on IGF1R, we employed lectin pulldown, treatment with glycosidases, and mass spectrometry-based glycoproteomics. We focused on the alpha subunit of IGF1R, as the binding site of ganitumab is located within this subunit^19^. Whole cell lysates were digested with sialidases that specifically cleave α2-3-linked sialic acids or broadly cleave α2-3,6 or α2-3,6,8-linked sialic acids, as specific sialidases for α2-6 or α2-8-linkages are not available. Treatment with α2-3 sialidase did not result in any apparent band shift compared to control, whereas α2-3,6 and α2-3,6,8 sialidases induced similar extent of IGF1R alpha subunit band shift, suggesting that the majority of sialic acids on IGF1R alpha are α2-6-linked **(Figure 2a)**. Next, we sought to determine if sialic acids on IGF1R are on N- or O-linked glycans. Treatment with peptide-*N*-glycosidase F (PNGase F), which digests all N-glycans, led to a significant band shift, showing a decrease of approximately 20 kDa. This is consistent with IGF1R being heavily N-glycosylated. No additional band shift was observed when PNGase F was combined with STI treatment **(Figure 2b)**, and PNGase F treatment completely abolished the ability of SNA to pull down IGF1R alpha **(Figure 2c)**, indicating that α2-6 sialic acids are predominantly present on N-glycans. In addition, treatment with STI and α2-3,6,8 sialidase abolished the binding of SNA to IGF1R **(Figures 2d and 2e)**. In the SNA blot, a band was observed around 100 kDa, indicating that IGF1R beta is also α2-6-sialylated **(Figure 2e and Supplementary Figures S3a and S3b)**. Notably, the pulldown of IGF1R alpha by SNA was also detected in lung, breast, and colorectal cancer cell lines, suggesting that α2-6 sialylation of IGF1R is present in these cell lines as well **(Figure 2f)**. Conversely, in normal human ovarian surface epithelial cells, no IGF1R alpha was identified in the SNA pulldown eluate, suggesting that the modification is specific to cancer cells **(Figure 2f)**. To pinpoint the specific asparagine (N) residues of IGF1R carrying sialyated glycans, a mass spectrometry (MS)-based glycoproteomic approach was employed. Analysis of the MS profile unveiled that the amino acid residue N607 of IGF1R was modified with sialylated glycans **(Figures 2g and 2h)**. Two types of sialylated glycans were found at N607, one complex biantennary N-glycan (HexNAc_4_Hex_5_NeuAc) and a hybrid N-glycan (HexNAc_3_Hex_5_NeuAc) **(Figure 2h and Supplementary Figure S3c).**

**Figure 2.**
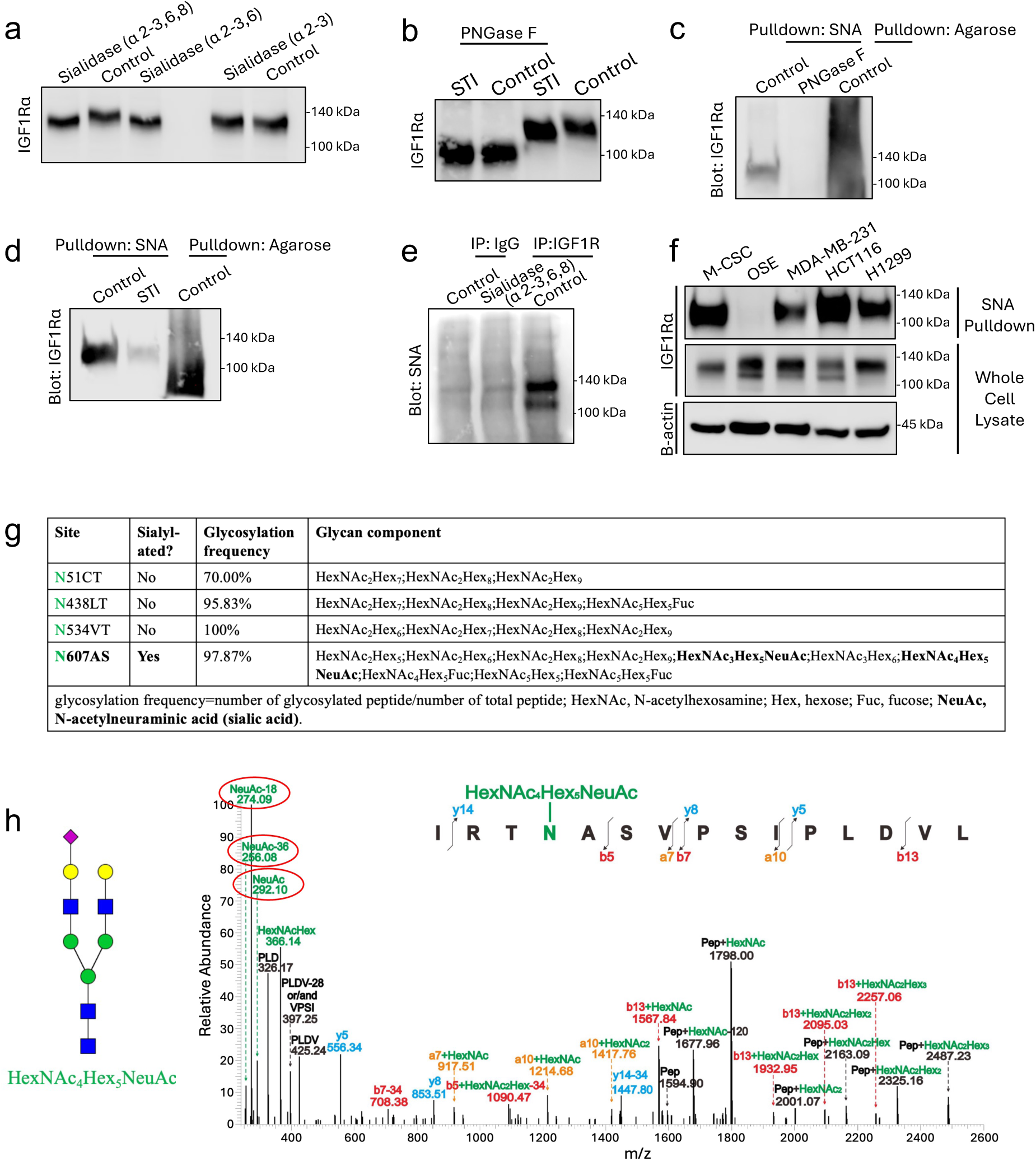
IGF1R alpha is modified with α2-6-linked sialylated N-glycans on asparagine 607. **(a)** Western blot of sialidase-treated IGF1R alpha **(b)** Western Blot of IGF1R alpha treated with STI alone or in combination with PNGase F. **(c)** Western blot of IGF1Rα after PNGaseF treatment and SNA pulldown. **(d)** Western blot of IGF1Rα after STI treatment and SNA pulldown **(e)** SNA lectin blot of sialidase-treated, immunoprecipitated IGF1R **(f)** Western blot of IGF1Rα after SNA pull down in ovarian cancer (SKOV-3), ovarian surface epithelial (OSE), breast cancer (MDA-MB-231), lung (HCT116), and colon cancer (H1299) cells. **(g)** Site-specific N-glycans of IGF1R alpha identified by mass-spectrometry (MS). **(h)** MS profile and annotation of identified sialylated glycan on Asn-607.

### ST6GAL1-mediated sialylation of IGF1R inhibits ganitumab binding in ovarian cancer patients

The human genome encodes two sialyltransferase paralogs, ST6GAL1 and ST6GAL2, which can add α2-6-linked sialic acids to N-glycans^20^. Through CRISPR/Cas9 knockout and SNA pulldown experiments, we determined that ST6GAL1 is the enzyme primarily responsible for sialylating IGF1R, while ST6GAL2 does not contribute to the process **(Figure 3a and Supplementary Figures S4a and S4b)**. Consistently, ST6GAL1 knockout significantly increased both immunoprecipitation and inhibition of IGF1R by ganitumab compared to cells transduced with non-targeting sgRNA **(Figures 3b and 3c)**. ST6GAL1 expression was found to be significantly upregulated in ovarian cancer tumor tissues compared to normal ovary in the cancer genome atlas (TCGA) database using UCSC Xenabrowser^21^ **(Figure 3d)**. Similarly, upregulation of ST6GAL1 was also observed in breast, colon, and lung cancers **(Supplementary Figure S4c)**. Next, we assessed if ST6GAL1 expression could be used as a determinant of the likelihood that a tumor tissue will exhibit enhanced ganitumab binding upon desialylation. To this end, ovarian cancer tumors were stratified into High and Low groups based on their ST6GAL1 H-scores **(Supplementary Figure S4d)**, and IHC was performed with Ganitumab after sialidase digestion. Ganitumab binding was more enhanced in ST6GAL1-High tumors compared to ST6GAL1-Low ones, suggesting that ST6GAL1-mediated sialylation was responsible for the dampened IGF1R-ganitumab interaction in these tissues. Since the variability of IGF1R expression between tumors is high, we also performed a correlation analysis between ST6GAL1 expression vs ganitumab signal normalized to total IGF1R expression. Our analysis revealed a significant negative correlation between ganitumab and ST6GAL1 signal intensities **(Supplementary Figure S4e)**, indicating a close clinical association.

**Figure 3.**
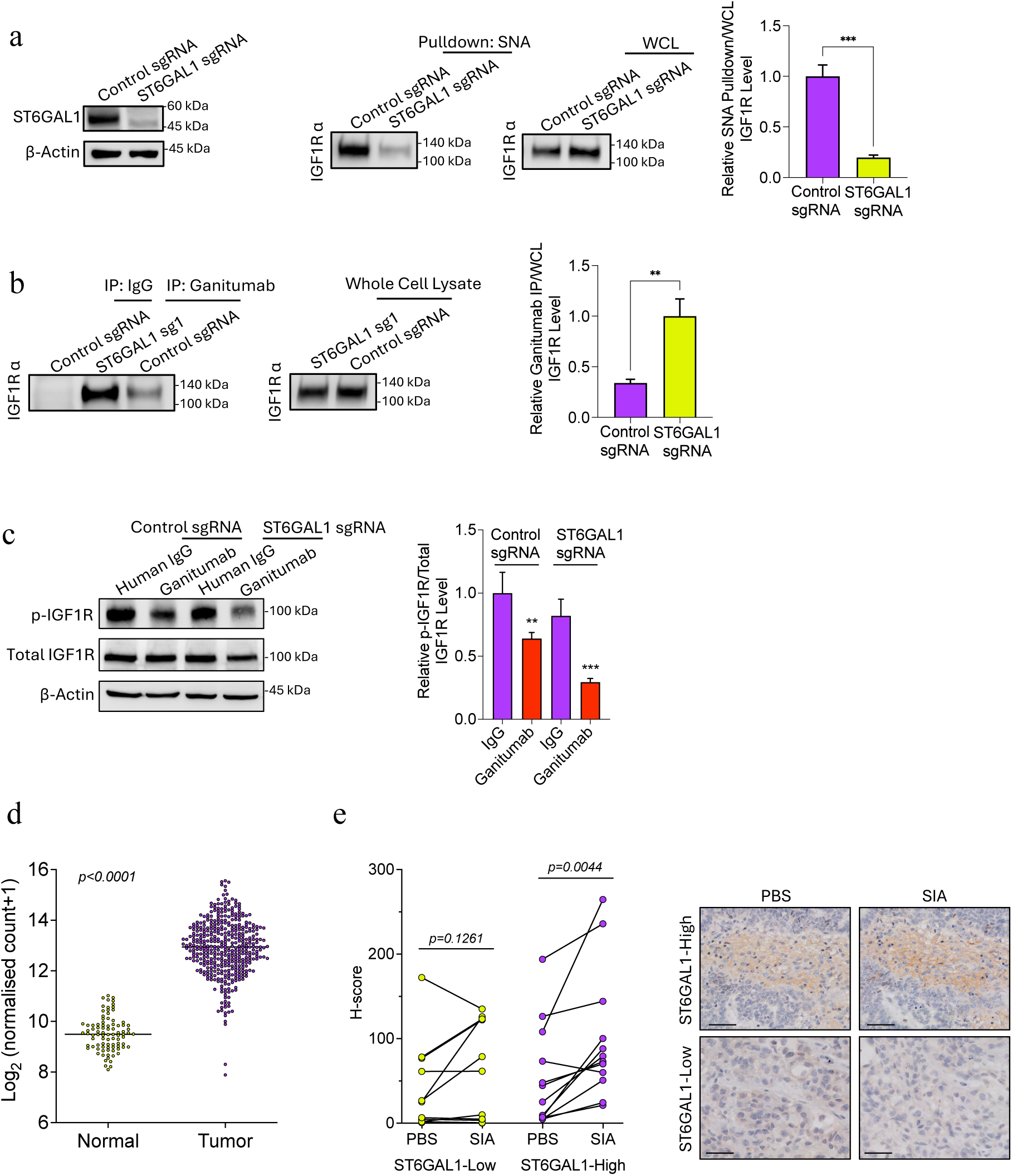
ST6GAL1-mediated IGF1R sialylation inhibits ganitumab binding in ovarian cancer patients. **(a)** Western Blot of IGF1R alpha after SNA pulldown in ST6GAL1 knockout cells **(b)** Western blot of IGF1R alpha immunoprecipitated with ganitumab in ST6GAL1 knockout cells **(c)** Western blot of p-IGF1R in ST6GAL1 knockout cells treated with ganitumab **(d)** ST6GAL1 expression in ovarian cancer tumor vs normal ovary tissue from TCGA data analyzed by Xenabrowser. **(e)** Ganitumab H-score of ovarian cancer FFPE tissues stratified into ST6GAL1-High (n=12) and ST6GAL1-Low (n=12) after sialidase (SIA) digestion; representative image of the staining and H&E staining of the tissues shown at 40x magnification. Scale bar = 50μm.

### N607 sialylation of IGF1R inhibits binding and efficacy of ganitumab

Next, we sought to evaluate the mechanism and role of N607 glycosylation in inhibiting IGF1R-ganitumab interaction. We utilized AlphaFold3^22^ to predict the structure of IGF1R-ganitumab complex **(Figure 4a)**. In this structure, N607 is situated at the junction loop between the FnIII-2 and FnIII-1 domains of IGF1R. We also conducted molecular dynamics (MD) simulations on the IGF1R-ganitumab structures with IGF1R’s N607 added with a sialylated glycan (HexNAc_4_Hex_5_NeuAc) or without it, and compared their RMSD and potential energy profiles to evaluate their stability **(Figure 4b)**. We found that the RMSD values of the glycan-modified IGF1R-ganitumab were significantly higher than those of the unmodified counterpart, suggesting that the glycosylation of IGF1R at N607 enhances the flexibility of the complex and likely destabilizes it. Next, we engineered an IGF1R mutant deficient in N607 glycosylation. To eliminate the effect of endogenous IGF1R, we generated IGF1R knockout SKOV-3 cells by CRISPR/Cas9 and introduced an IGF1R variant carrying an asparagine to glutamine (Q) mutation at amino acid residue 607 (IGF1R N607Q). In comparison to wildtype IGF1R (IGF1R WT), IGF1R N607Q exhibited a band shift of IGF1Rα in Western Blot **(Figure 4c)**, indicating that amino acid residue 607 is defective in glycosylation. Furthermore, IGF1R N607Q showed significantly lower sialylation levels compared to WT, as demonstrated by Western Blot following SNA pulldown **(Figure 4d)**. Importantly, ganitumab immunoprecipitation and inhibition of IGF1R phosphorylation was notably enhanced with IGF1R N607Q **(Figure 4e and Supplementary Figure S5a)**, suggesting that α2-6-sialylation of IGF1R N607 impedes ganitumab binding and subsequent deactivation of IGF1R. In addition, the IGF1R N607Q mutant displayed significantly fewer colonies under ganitumab treatment in clonogenic cell survival assay, indicating that deglycosylation of N607 is sufficient to sensitize ovarian cancer cells to ganitumab **(Figure 3f)**. Although levels of p-IGF1R and total IGF1R levels were reduced by ganitumab in IGF1R WT-expressing cells **(Supplementary Figure S5a)**, colony formation remained unaffected. This observation suggests that the expression level is unlikely to be an indicator of ganitumab response and supports our proposition that IGF1R sialylation acts as a resistance mechanism. Subsequently, we investigated the *in vivo* impact of N607 deglycosylation on sensitizing ovarian cancer cells to ganitumab. IGF1R knockout ovarian cancer cells expressing either IGF1R WT or the IGF1R N607Q mutant were intraperitoneally injected into nude mice and treated with ganitumab. Bioluminescence imaging revealed that ganitumab treatment did not significantly affect the control group with WT IGF1R, whereas a substantial suppression of metastatic growth was observed in the group with IGF1R N607Q mutation following ganitumab treatment **(Figure 3g)**. Similar differences were observed in the number of tumor nodules on the mesenteries (**Figure 3g**). Tumors with the IGF1R N607Q mutation exhibited metastatic growth comparable to those with wildtype IGF1R **(Figure 3g),** indicating that N607 deglycosylation alone did not impact the oncogenic properties of IGF1R. This is consistent with our observation that desialylation alone has no effect on the activity of IGF1R **(Supplementary Figure S2c)**. IHC staining of omental metastatic tumors from these mice showed significantly lower phospho-IGF1R signals, as determined by H-score, indicating enhanced deactivation of IGF1R due to ganitumab treatment **(Supplementary Figure S5b).** Taken together, these results suggest that deglycosylation of asparagine 607 of IGF1R can effectively overcome ganitumab resistance.

**Figure 4.**
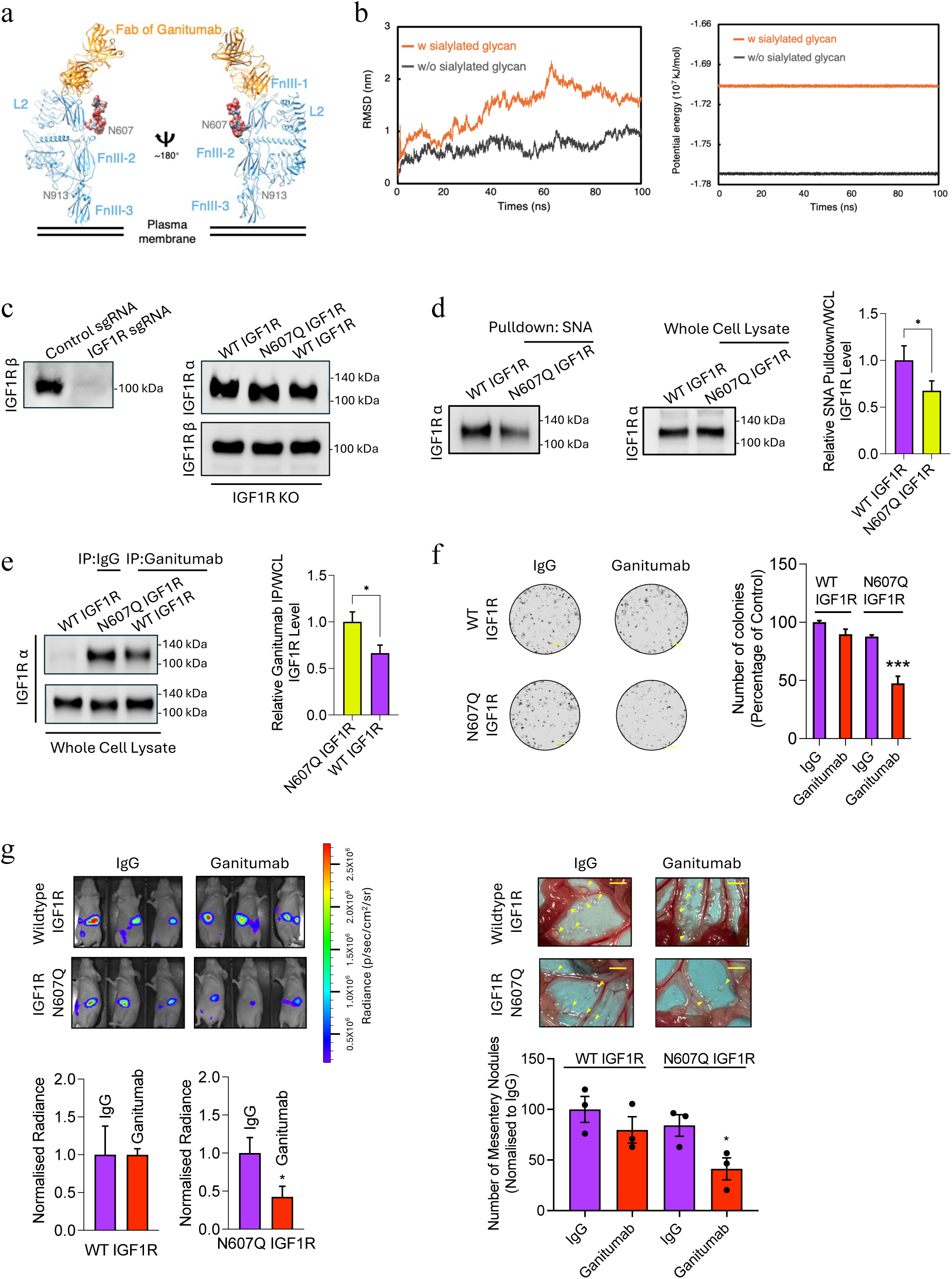
Asparagine 607 sialylation of IGF1R inhibits binding and response to ganitumab. **(a)** Structure of predicted IGF1R-ganitumab complex. The HexNAc4Hex5NeuAc group is displayed in surface mode. (**b**) RMSD profiles and potential energy from the MD simulation results of IGF1R-ganitumab with IGF1R’s N607 modified either with (orange line) or without (gray) a sialylated glycan. **(c)** Western blot of IGF1R alpha and beta subunits in wildtype (WT) and N607Q mutant. **(d)** Western blot of IGF1R N607Q after SNA pulldown **(e)** Western Blot of IGF1R N607Q alpha immunoprecipitated with ganitumab. **(f)** Clonogenic cell survival assay of IGF1R N607Q treated with ganitumab; scale bar= 0.5mm. **(G)** Bioluminescence imaging and mesenteric tumor nodules of ganitumab-treated ovarian cancer cells harboring N607Q IGF1R mutation; n=4 per group; scale bar= 0.25cm.

## Discussion

IGF1R is a heavily sought-after drug target, but attempts to develop an efficacious agent against this receptor have been unsuccessful in oncology. Numerous studies have delved into the resistance mechanisms of anti-IGF1R therapeutics^1,23,24^. These include constitutive activation of downstream signaling pathways^25,26^ and compensation by other receptors^27,28^. However, despite attempts to enhance treatment outcome through combination trials with other agents, success has been limited^29^, indicating the need to explore alternative strategies for enhancing the efficacy of anti-IGF1R therapies. One promising yet often overlooked area is the impact of intrinsic resistance due to post-translational modifications. This oversight may stem from insufficient insight into how these modifications can affect protein behaviour and binding dynamics. In this study, we have uncovered a tumor-specific post-translational N-glycan sialylation at a specific residue of IGF1R that is prevalent in various cancer types, offering a novel mechanistic explanation for the observed resistance.

Altering just a single sugar residue can significantly impact the function of a protein. The presence of glycans can affect binding affinities and overall dynamics of protein complexes^16^. Here we found that sialylation of IGF1R amino acid residue N607 impacts ganitumab binding on the alpha subunit of IGF1R. Interestingly, this binding site is not located at the predicted interface of the ganitumab-IGF1R co-complex structure. With our N607 IGF1R site-specific mutant, we observed enhanced ganitumab efficacy over wildtype IGF1R *in vivo*, confirming a specific role for N607. Our MD analysis comparing the RMSD profiles of the IGF1R-ganitumab complex with and without the sialylated glycan at N607 revealed that this modification destabilizes the complex. These findings are consistent with a recent study on SARS-CoV-2, which demonstrated that O-glycans help in maintaining interactions with the host receptor by forming polar contacts with nearby residues without disrupting the direct binding site^30^. Moreover, we observed no change in IGF1R protein levels, activity, or cell surface localization after desialylation, implying that this modification may not affect the receptor’s overall structural integrity or membrane availability. Indeed, research has indicated that N-glycans can be dynamic and accessible, influencing protein interactions without necessarily altering the overall stability of the protein structure^31^. Glycosylation of the IgG domain of antibodies is known to determine their ability to interact with antigens and is an active area of research. However, much less is known about how the glycosylation of proteins affects antibody binding^32^. Our findings further underscore the importance of protein glycan modifications in the design of therapeutic antibodies. As mAbs and antibody conjugates continue to dominate the biopharmaceutical landscape, integrating glycan engineering into the antibody screening process will be essential for optimizing targeting accuracy and efficiency.

Identifying specific glyco-modifications linked to distinct phenotypes remain challenging due to the vast diversity in glycans and various technical constraints. The majority of current studies investigating the connection between glycosylation and therapy response are carried out *in vitro* using cell or biochemical assays, lacking animal models and direct mechanistic insight^14,33–35^. We utilized a combination of mouse model, patient sample and biochemical analysis to precisely map the site and regulation of the glyco-modification responsible for this resistance. We identified N607 as the pivotal amino acid residue where sialylated N-glycans mediate resistance to ganitumab. Importantly, these findings were supported by protein structure analysis and *in vivo* experiments. Our detailed findings are crucial for translational applications, as sialic acids have essential physiological functions^17^ and their global inhibition could lead to undesirable side effects. This may explain why sialyltransferase inhibitors are not in clinical trials, despite several studies demonstrating their effectiveness as anti-cancer agents^36^. While N-glycans comprise a complex array of glycans with numerous essential functions that cannot be broadly targeted, sialylation represents a terminal modification that can be selectively targeted. Moreover, the loss of glycosylation at the specific amino acid residue (N607) identified in this study has no discernable effect on protein maturity and function, further supporting the basis for specifically targeting it. Future therapeutic strategies could therefore benefit from targeting sialylated IGF1R, potentially through methods such as development of glycoform-specific antibodies or antibody-sialidase conjugates^37^.

The inability to identify patients most suited for anti-IGF1R therapy is another significant factor contributing to the failure of these trials^24^. Our identification of ST6AL1 as the key enzyme responsible for IGF1R sialylation can provide crucial insights for patient stratification. ST6GAL1 is upregulated in several solid tumors^38^, and serves as the primary sialylatransferase responsible for adding α2-6 sialic acids to N-glycans on many cell surface receptors. Various studies have suggested ST6GAL1 could induce therapeutic resistance, but information regarding its impact on patients is limited^15,34,35^. Duarte et al. demonstrated the presence of α2-6 sialic acid, the product of ST6GAL1-mediated sialylation, on ErbB2 receptors in gastric cancer patient samples^15^; however, direct experimental evidence linking ST6GAL1 to anti-ErbB2 mAb-tumor binding is lacking. Kim et al. observed that IGF1R N913 was glycosylated in some figitumumab-sensitive cancer cell lines^39^, however experiments were performed with deglycosylated IGF1R in a sensitive cell line, making it difficult to predict if altering this glycan would help in overcoming resistance. In other cases where a more direct benefit of desialylation is evident, it includes additional factors such as the involvement of the immune system^40^. This makes it challenging to predict therapy response in patients. In contrast to these prior studies which have primarily yielded indirect evidence or mechanism, we are the first to employ enzymatic antigen retrieval of FFPE patient samples to demonstrate a direct connection between ST6GAL1, sialylation, and the binding of a therapeutic mAb. In clinical settings were glycoproteomic data is available, the sialylation status of IGF1R N607 and expression of ST6GAL1 could be utilized to select patients most likely to benefit from anti-IGF1R mAb-based therapies.

In conclusion, our study not only elucidated the critical site of glycosylation but also demonstrated that removal of the sialylated glycan can effectively alter resistance to anti-IGF1R therapy. The presence of this modification in several other cancers, including breast, colon, and lung cancers, but not normal tissues, implies that our findings may have broad implications, potentially explaining the lack of success of ganitumab in clinical settings for multiple cancers. Moreover, our data also suggests that the expression of ST6GAL1 and IGF1R N607 sialylation status could serve as a marker to stratify patients likely to respond to anti-IGF1R therapy, which may benefit cancers that exhibit resistance to therapy due to hypersialylation.

## Materials and Methods

### Cell lines and cell culture

Cancer cell lines were cultured in a humidified CO_2_ (5%) incubator at 37 °C. Ovarian cancer cell line SKOV-3 and normal human ovarian surface epithelial (OSE) cells, gifts from Dr. N. Auersperg (University of British Columbia, Canada), were cultured in Medium 199 and MCDB105 in 1:1 ratio. MDA-MB-231 (breast), HCT116 (colon), and H1299 (lung) cancer cells were maintained in RPMI-1640 medium. SKOV3 and OSE cells were supplemented with 5% fetal bovine serum (FBS) and 15% FBS, respectively, and 1% penicillin and streptomycin (100 units/ml). All other cancer lines were supplemented with 10% FBS and 1% penicillin and streptomycin.

### Patient tumor samples

Ovarian cancer samples were obtained from patients who underwent surgery at Queen Mary Hospital, University of Hong Kong with their written consent. The samples were fixed in formalin and embedded in paraffin. Histological sections from these samples were stained as described below under **Immunohistochemistry and lectin histochemistry**. Clinicopathological features of the patients are included in **Table S1**.

### Western blot and lectin blot

Cells were lysed in buffer containing 62.5 mM Tris-Cl and 2% sodium diodecyl sulfate (SDS). Proteins were denatured by boiling in 0.1 M dithiothreitol (DTT) before being subjected to sodium dodecyl sulphate-polyacrylamide gel electrophoresis SDS-PAGE. Next, proteins were transferred to nitrocellulose membrane (BioRad) and blocked in 5% non-fat milk in phosphate buffered saline with 0.05% Tween-20 (PBS-T). Primary antibody or biotinylated lectin incubation was performed overnight followed by incubation with appropriate horseradish-conjugated secondary antibody. The antibodies and lectin used include: IGF1R α (Santa Cruz, Catalog# sc-271606), IGF1R β (Cell Signaling Technology, Catalog# 3027), phospho-IGF1R β (Cell Signaling Technology, Catalog# 4568), ST6GAL1 (R&D Systems, Catalog# AF594), β-actin (Thermo Fisher, Catalog# MA1-140), and *Sambucus Nigra* (SNA) lectin (Vector laboratories, Catalog# B-1305-2).

For Western/lectin blot experiments involving glycosidase digestion, cells were lysed in radioimmunoprecipitation assay (RIPA) buffer and whole cell protein lysates were subjected to glycosidase digestion (1h for sialidase and fucosidase; overnight for PNGaseF) at 37 °C, following by denaturation and SDS-PAGE. Glycosidases used include: α 2-3 sialidase (Sigma, Catalog# N7271), α 2-3,6 sialidase (Sigma, Catalog# N5521), α 2-3,6,8 sialidase (New England Biolabs, Catalog# P0720), α 1-3,4 Fucosidase (New England Biolabs, Catalog# P0769), and PNGase F (New England Biolabs, Catalog# P0704).

### Immunoprecipitation and lectin pulldown

Cells were lysed in radioimmunoprecipitation assay (RIPA) buffer and incubated at 4 °C in an end-to-end shaker. Cell debris was removed by centrifugation at 14,000g for 10 minutes. 1 μg total protein was incubated with anti-IGF1R antibody or agarose-bound lectin overnight. Protein A magnetic beads (Cell Signaling Technology, Catalog# 73778) were used to precipitate antibody-protein complex. For elution of protein from antibody-protein complex and lectin-protein complex, manufacturer’s protocol was followed. In experiments involving glycosidase digestion, it was performed prior to addition of antibody/lectin.

### Mass spectrometry (MS)-based glycoproteomics

IGF1R was immunoprecipitated as described above and subjected to SDS-PAGE. The gel was stained, and then destained as described elsewhere^40^. The gel bands corresponding to IGF1R alpha and beta subunits (based on molecular weight) were excised and then treated as described elsewhere ^41^. Briefly, the sample was reduced using DTT, carbamidomethylated using iodoacetamide, and then treated with either MS-grade trypsin (Thermo Scientific, Catalog# 90057) or chymotrypsin (Thermo Scientific, Catalog# 90056). The enzyme digested protein was purified using Pierce™ C18 tips (Thermo Scientific, Catalog# 87782) and then subjected to liquid chromatography-mass spectrometry (LC-MS) using Q Exactive Plus with an EASY-spray source and EASY-nLC 1200 system (Thermo Fisher Scientific) for identification of N-glycopeptides. The instrument was operated in a data-dependent mode, the MS precursor scan was performed at resolution set to be 70,000 in positive mode, automatic gain control (AGC) target was set to be 1e^6^, and maximum injection time (IT) was 30 ms, MS/MS fragment ions were analyzed at resolution set to be 17,500, the AGC target was set to be 5e^4^, and maximum IT was 500 ms, the twenty most intense peptide ions (TOP 20) were sequentially isolated and analyzed using normalized collision energy (NCE) of 35%. The glycoproteomic MS raw data were searched by software Byonic (version 5.1) against the IGF1R fasta sequence (P08069) obtained from UniProt. Different search parameters were used for the protein identity validation, with carboxyamidomethylation of cysteine as a fixed modification, oxidation of methionine, deamidation of asparagine and glutamine as a variable modification. When trypsin was selected as the digestion enzyme, the carboxyl side of lysine and arginine was set as the cleavage site, and up to 2 missed cleavages were allowed. When chymotrypsin was selected as the digestion enzyme, the carboxyl side of tyrosine, phenylalanine, tryptophan, and leucine was set as the cleavage site, and up to 2 missed cleavages were allowed. The digestion specificity was set to be fully specific. The glycoproteomic data were subsequently manually processed using Xcalibur (version 4.2.28.14) to obtain the glycosylation sites and corresponding glycan components. The glycopeptides were further checked by manual examination of the corresponding MS/MS spectra via oxonium ions, monosaccharides, oligosaccharide neutral loss patterns. List of identified glycans is listed in **Table S2**.

### Structural prediction and molecular dynamics (MD) simulation

The structure of IGF1R-ganitumab complex was predicted using AlphaFold3 ^22^. In the structural prediction, we removed the intercellular regions of IGF1R and the Fc domain of ganitumab. For the MD simulation, the systems of IGF1R-ganitumab structures with or without a sialylated glycan were solvated using the TIP3P water model ^42^ and neutralized in a 150 mM NaCl environment. Each periodic system was contained in a cubic simulation box with approximate dimensions of 232.93×232.93×232.93 Å. The systems were parameterized using the CHARMM36m force field ^43^ and all MD simulations were performed with GROMACS ^44^. Energy minimization was carried out using the steepest descent method. Production runs were conducted under periodic boundary conditions in the NPT ensemble, employing a velocity-rescale thermostat and a Cell Rescaling barostat to maintain the temperature at 310 K and the pressure at 0.8 bar, respectively. The simulations were conducted for 100 ns with a time step of 2 fs.

### CRISPR/Cas9

SKOV-3 cells were stably transduced with Lenti-Cas9-Blast (Addgene, Plasmid# 52962). Cas9-expressing cells were selected with 10 µg/ml Blasticidin (Gibco, Catalog# A1113903). Guide RNAs were cloned into Lenti-Guide-Puro (Addgene Plasmid# 52963) and transduced in Cas-9 expressing cells with target sgRNA followed by 1 µg/ml Puromycin (Gibco, Catalog# A1113803) selection as previously described ^45^. Sequences of sgRNA used: 5’-GGAGAACGACCATATCCGTG-3’ (IGF1R), 5’-TGTGAATGTATCCATGGTAG-3’ (ST6GAL1), 5’-GGAGAACGACCATATCCGTG-3’ (ST6GAL2), 5’-AGCACGTAATGTCCGTGGAT-3’ (Non-specific sgRNA). Efficiency of sgRNA was evaluated using TIDE (Tracking of Indels by DEcomposition) ^46^ and Western Blot.

### Plasmid construction for IGF1R mutation

Mutation of IGF1R asparagine 607 to glutamine (N607Q) was performed using overlap PCR with wildtype IGF1R as template. Briefly, 10 cycles of PCR were performed using the following primers: forward primer, 5’-ATCTTGTACATTCGCACC**CAG**GCTTCAGTTCCTTCCATT-3’ and reverse primer: 5’-AATGGAAGGAACTGAAGC**CTG**GGTGCGAATGTACAAGAT-3’, followed by PCR product purification. The purified product was then used as template in 25 cycles of PCR using full-length-IGF1R-amplifying primers with restriction enzyme (RE) sites (forward primer, 5’-CCTCTAGAGCCACCATGAAGTCTGGCTCCGGAGG-3’) and reverse primer, 5’-CCGGATCCAGCGTAATCTGGAACATCGTATGGGTAGCAGGTCGAAGACTG-3’). The mutant IGF1R construct was then RE-digested, cloned into a lentiviral vector, and stably transduced in SKOV-3 IGF1R knockout cells.

### Clonogenic cell survival assay

Ovarian cancer cells were pre-treated with 1 µg/ml ganitumab (MedchemExpress, Catalog # HY-P99294), 200 µM sialyltransferase inhibitor (STI; 3Fax-Peracetyl Neu5Ac) (Sigma-Aldrich, Catalog# 566224), or 200 µM fucosyltransferase inhibitor (FTI; 2F-Peracetyl-Fucose) (Sigma-Aldrich, Catalog #344827) for 48 hours. 300 cells were then seeded in 6-well plates and cultured for 10 days under ganitumab, STI, or FTI treatment. Colonies were fixed in methanol and stained with 1% crystal violet for visualization and then counted.

### *In vivo* xenograft model of ovarian cancer metastasis

Nude mice (4-6 weeks old) were intraperitoneally injected with 3 × 10^6^ luciferase-expressing SKOV-3 cells. After 1 week, mice were intraperitoneally injected with 6 mg/kg ganitumab or 10 mg/kg STI three times a week for 2 weeks. At the endpoint of the experiment, animals were intraperitoneally injected with luciferase substrate and imaged using IVIS Spectrum (Perkin Elmer) to evaluate the extent of tumor growth. Animals were then sacrificed, and metastatic dissemination was evaluated by counting the number of tumor nodules on mesenteric membrane. Tumors that metastasized to the omentum were harvested and fixed in 4% formalin overnight and embedded in paraffin.

### Immunohistochemistry and lectin histochemistry

Paraffin-embedded tissue sections were hydrated in water after dewaxing in xylene and ethanol. Antigen retrieval was performed by boiling samples in sodium citrate buffer, followed by peroxidase block using 3% H_2_O_2_. In experiments where sialidase digestion was performed on IHC slides, tissue sections were incubated in 5% α2-3,6,8 neuraminidase (New England Biolabs, Catalog# P0720) in PBS overnight in a humidified chamber at 37°C prior to peroxidase blocking. For immunohistochemistry, staining was performed using IHC Staining Kit (Abcam, Catalog# ab64264) following manufacturer’s instructions. For lectin histochemistry, tissue sections were blocked with CarboFree Blocking Solution (Vector laboratories, Catalog# SP-5040-125) and stained with SNA lectin at 1: 4000 dilution following manufacturer’s instructions. In experiments where ganitumab signal intensity was quantified, slides were scanned with Vectra Polaris (Akoya Biosciences) and analyzed using QuPath software ^47^. For cell blocks, the mean DAB signal intensity per cell (mean cell DAB OD) in treatment group was normalized that of the respective control group to obtain the relative mean cell DAB OD. For H-scoring, arbitrary signal threshold was set to identify negative, weak staining, moderate staining, and strong staining in QuPath as previously reported ^48^. The H-score was computed by multiplying weak, moderate, and strong staining by a factor of 1, 2, and 3 respectively.

### Statistical analysis

Statistical analysis was performed using GraphPad Prism. Statistical significance between two independent means was evaluated using Student’s t-test for data that follow normal distribution. In other cases, Mann-Whitney U test was used. Pairwise t-test was used to compare ganitumab staining intensity in the same patient section with and without sialidase digestion. For multiple means, one-way ANOVA and post hoc Dunnet test was used. Error bars represent standard deviation, except in figures involving animal experiments, where it represents standard error of mean (SEM). A p value of less than 0.05 was considered statistically significant. *= p<0.05, **= p<0.005, *** =p<0.0005

## Acknowledgements

We acknowledge the funding support from “Laboratory for Synthetic Chemistry and Chemical Biology” under the Health@InnoHK Program launched by Innovation and Technology Commission, HKSAR, the Natural Science Foundation of China (22177097), and the Research Grants Council Senior Research Fellow Scheme (SRFS2223-7S05 to A. S. T. Wong and SRFS2324-7S01 to X. Li) and General Research Fund (17104022 to L. W. T. Cheung).

## Author Contributions

A.A.H. designed and performed experiments, analysed results, and prepared the manuscript. Q.C. designed and performed mass-spectrometry-based glycoproteomic analysis and revised the manuscript. K.S.W.F. helped with mice experiments and staining and analysis of patient samples. S.K.Y.T. and L.W.T.C. helped with revision and preparation of manuscript. C.Z. and N.L. performed structural prediction and MD simulation. C.N.L. and C.M.C. provided cancer cell lines. P.P.C.I. provided patient samples. X.L. and A.S.T.W. designed and supervised the project. All authors reviewed the manuscript.

