## Supplementary Figures and Tables for "Cancer-specific sialylation of insulin-like growth factor 1 receptor impairs therapeutic antibody binding and efficacy"

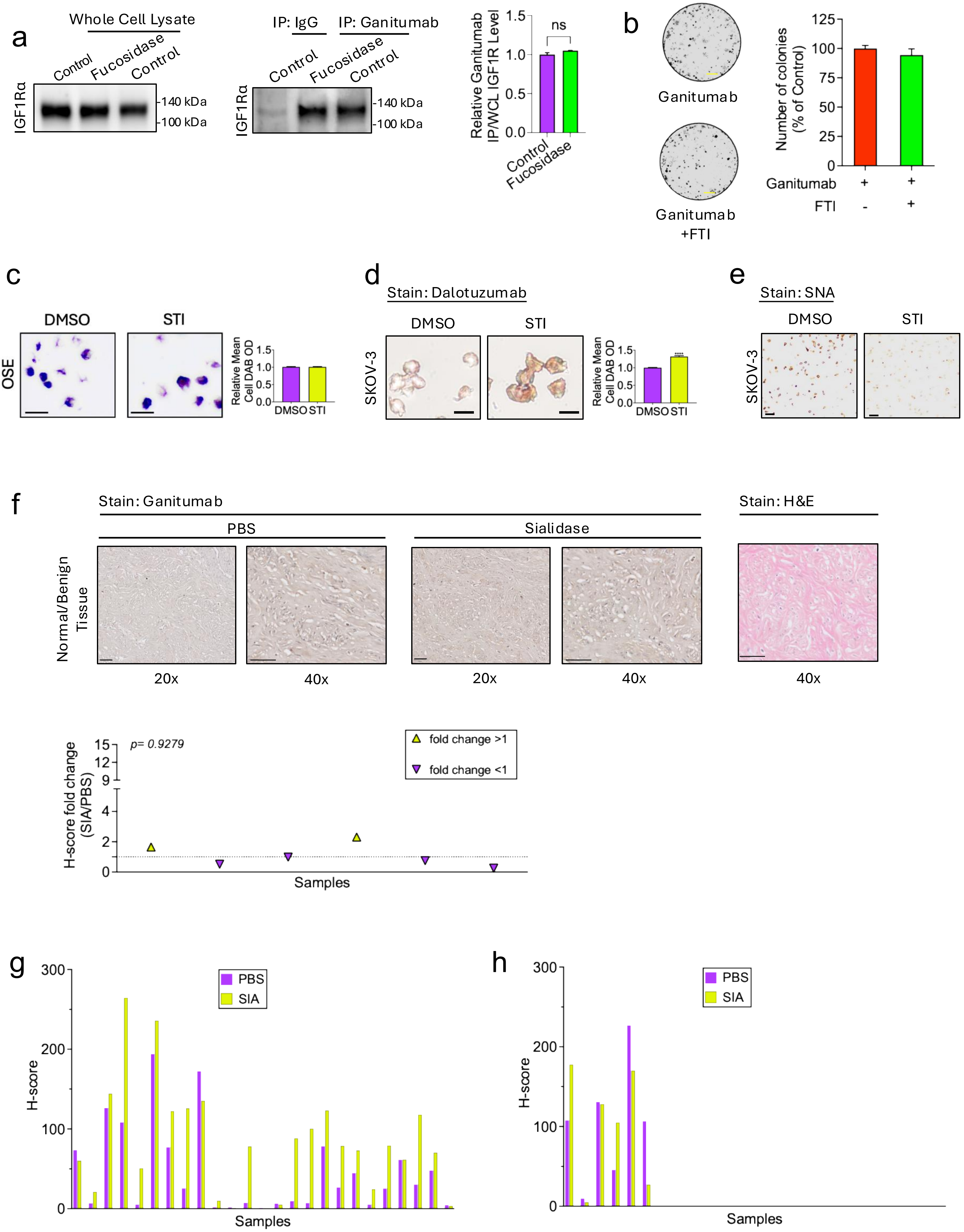
a
Whole Cell Lysate
Control
Fucosidase
Control
IGF1Rα
140 kDa
100 kDa
IP: IgG
IP: Ganitumab
Control
Fucosidase
Control
IGF1Rα
140 kDa
100 kDa
Relative Ganitumab IP/WCL IGF1R Level
ns
Control
Fucosidase
b
Ganitumab
Ganitumab +FTI
Number of colonies (% of Control)
Ganitumab + +
FTI - +
c
DMSO
STI
OSE
Relative Mean Cell DAB OD
DMSO STI
d
Stain: Dalotuzumab
DMSO
STI
SKOV-3
Relative Mean Cell DAB OD
DMSO STI
e
Stain: SNA
DMSO
STI
SKOV-3
f
Stain: Ganitumab
PBS
Sialidase
Stain: H&E
Normal/Benign Tissue
20x
40x
20x
40x
40x
p= 0.9279
fold change >1
fold change <1
H-score fold change (SIA/PBS)
Samples
g
PBS
SIA
H-score
Samples
h
PBS
SIA
H-score
Samples

**Figure S1. (a)** Immunoprecipitation of fucosidase-treated insulin-like growth factor 1 receptor (IGF1R) with ganitumab. **(b)** Clonogenic cell survival assay of ovarian cancer cells treated with ganitumab only or in combination with fucosidase (FTI) ; scale bar= 0.5mm. **(c)** Ganitumab staining of FFPE section from normal epithelial ovarian cell block (OSE) treated with STI; scale bar = 25µm. **(d)** Dalotuzumab staining of STI treated SKOV-3 cell block; scale bar = 25µm. **(e)** SNA staining of formalin-fixed, paraffin-embedded (FFPE) section from ovarian cancer cell block (SKOV-3) treated with STI. Scale bar = 100µm. **(f)** Ganitumab H-score fold change of normal/benign gynecological (n=6) FFPE tissues after sialidase (SIA) digestion; representative images of the staining and H&E staining of the tissues were shown at 20x and 40x magnification. Scale bar = 50µm. **(g)** Tumor tissue and **(h)** normal/benign tissue ganitumab H-scores after SIA digestion.

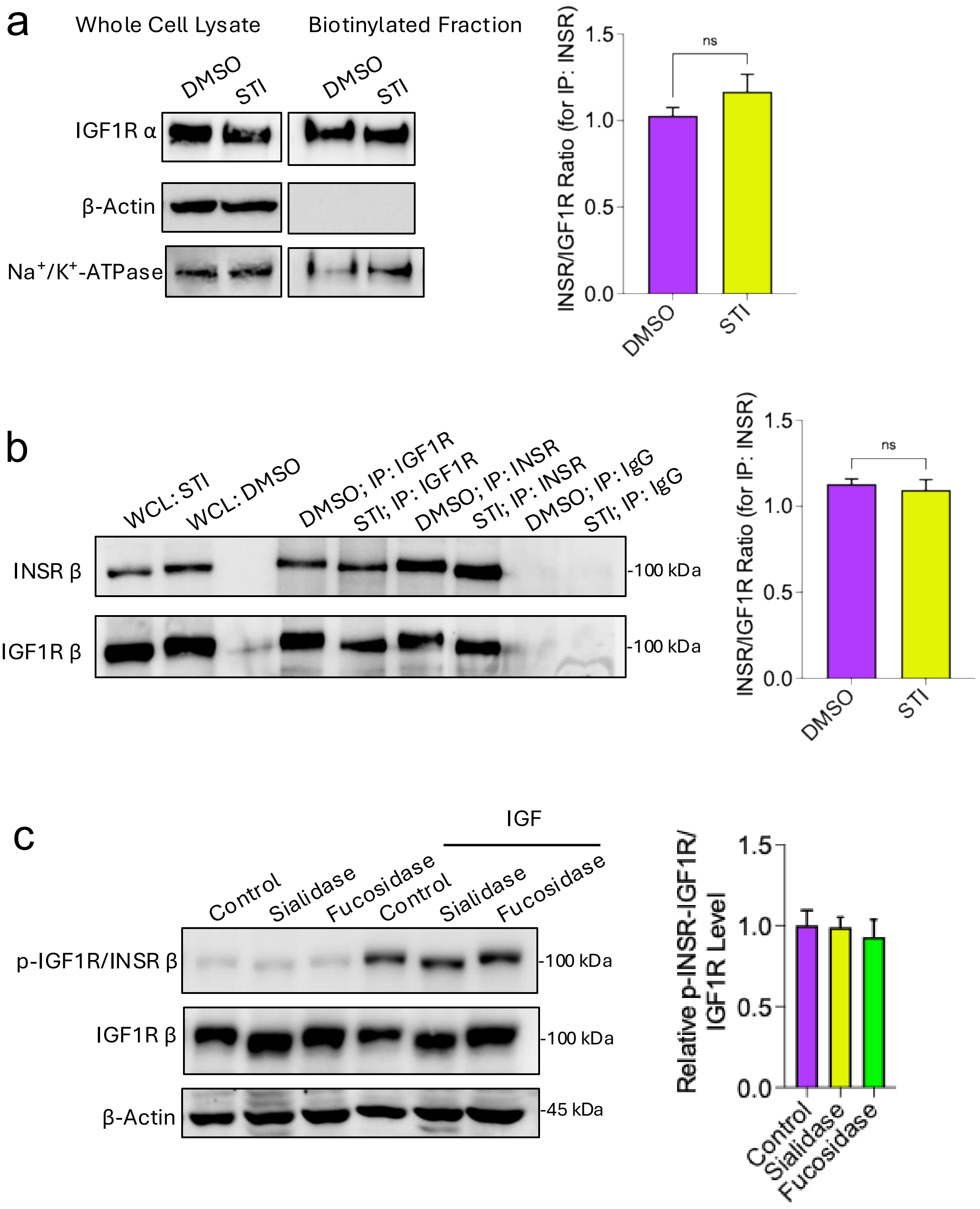


**Figure S2.** **(a)** Western blot of IGF1R alpha biotinylated cell-surface fraction. **(b)** Western blot of immunoprecipitated IGF1R beta and INSR beta after STI treatment. **(C)** Western Blot of p-IGF1R beta/INSR beta treated with sialidase or fucosidase

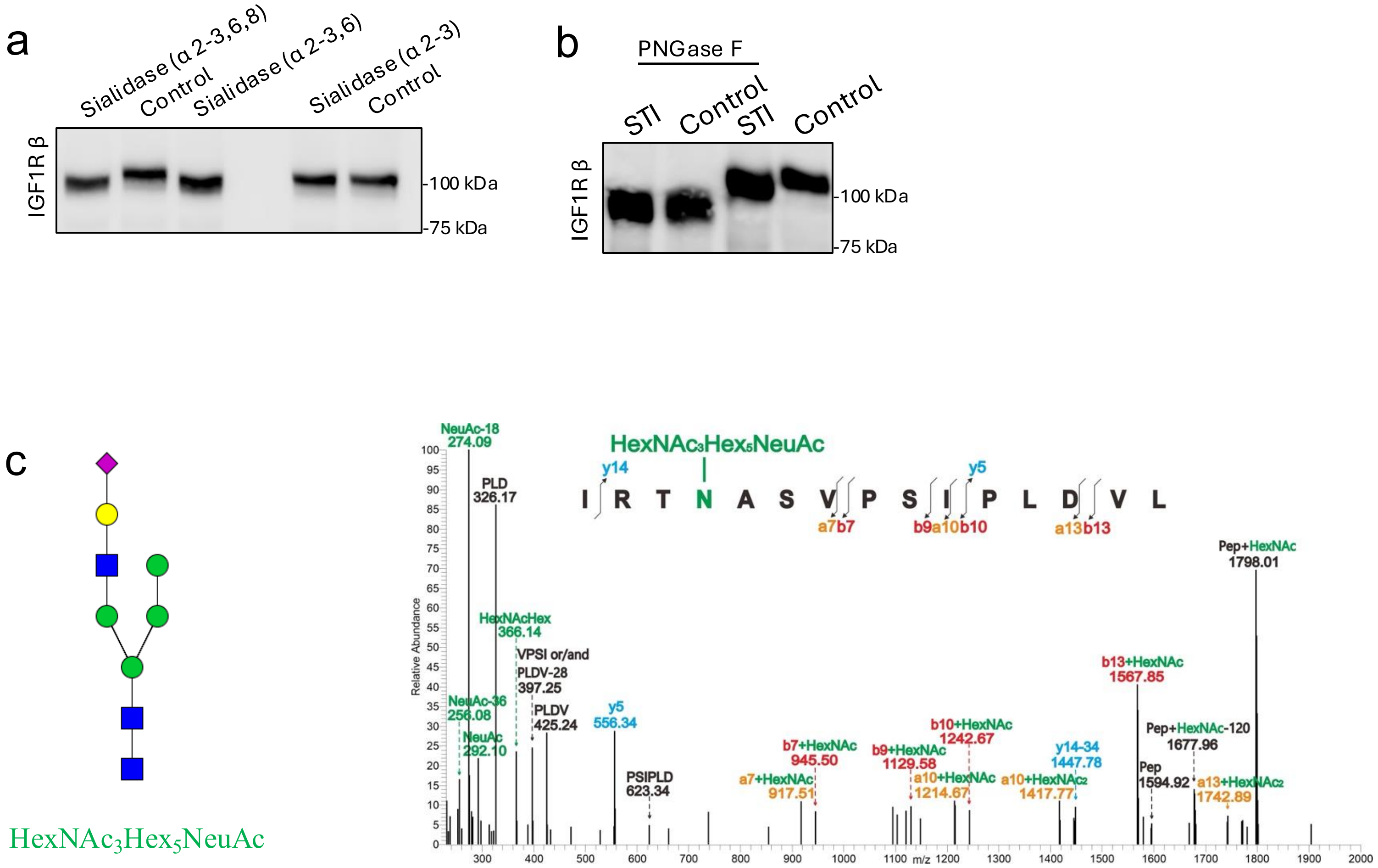


**Figure S3. (a)** Western blot of sialidase-treated IGF1R beta. **(b)** Western Blot of IGF1R beta treated with STI alone or in combination with PNGase F. **(c)** Mass-Spectrometry profile annotation of hybrid sialylated N-glycan at N607

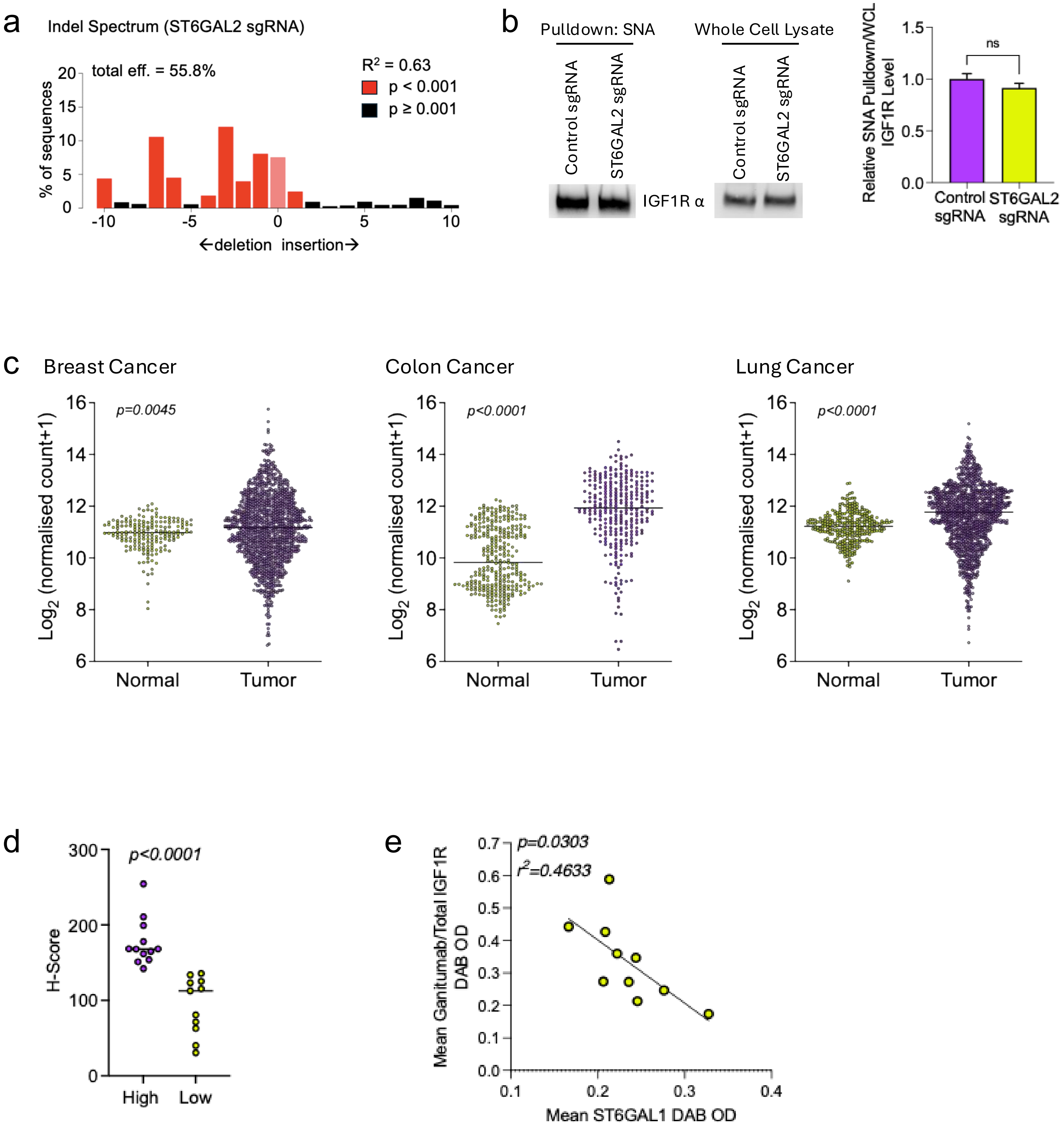


**Figure S4**. **(a)** Indel spectrum of ST6GAL2 sgRNA-transduced cells assessed by Tracking of Indels by DEcomposition (TIDE). **(b)** Western blot of IGF1R from ST6GAL2 sgRNA-transduced cells after SNA pulldown. **(c)** ST6GAL1 expression in breast, colon, and lung cancer tumor vs matched normal tissue from TCGA data analyzed by Xenabrowser. **(d)** ST6GAL1 H-score of ovarian cancer patient tumors used for stratification in **Figure 3e**. **(e)** Correlation of ST6Gal I and Ganitumab (normalized to total IGF1R) signal from IHC in ovarian cancer patient samples.

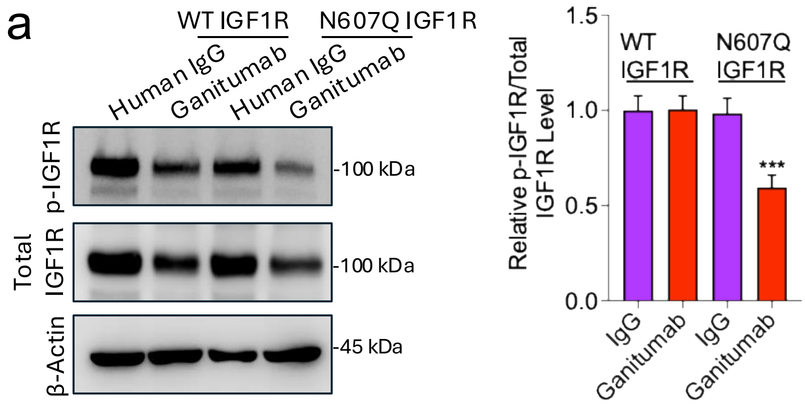


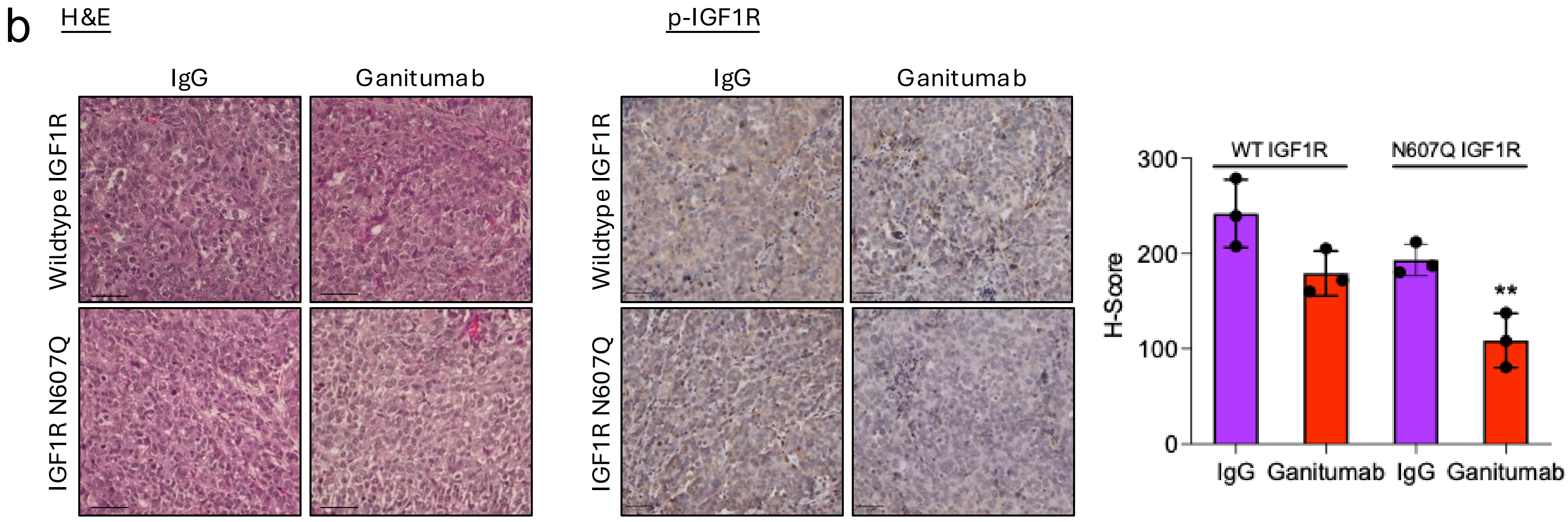


**Figure S5. (a)**Western blot of p-IGF1R N607Q beta after treatment with ganitumab. **(b)** IHC staining of p-IGF1R and H&E of ganitumab-treated xenograft tumors harvested from mice in **Figure 4g**; scale bar = 50μm.

| **Patients, N** | 25 |
|---|---|
| **Age (years)** | |
| Range | 35-70 |
| Mean ± SEM | 48.5 ± 1.7 |
| **Stage** | |
| I-IIA | 16 |
| IIB-IV | 9 |
| **Histological Subtype** | |
| High-grade serous carcinoma (HGSC) | 17 |
| Clear Cell Carcinoma (CCC) | 7 |
| Endometrial Carcinoma (EC) | 1 |

**Table S1. Clinicopathological table of patient samples used in the study**

| **Site** | **Sub-unit** | **Sialyl-ated?** | **Glycosylation frequency** | **Glycan component** |
|---|---|---|---|---|
| N51CT | α | No | 70.00% | HexNAc$_2$Hex$_7$;HexNAc$_2$Hex$_8$;HexNAc$_2$Hex$_9$ |
| N438LT | α | No | 95.83% | HexNAc$_2$Hex$_7$;HexNAc$_2$Hex$_8$;HexNAc$_2$Hex$_9$;HexNAc$_5$Hex$_5$Fuc |
| N534VT | α | No | 100% | HexNAc$_2$Hex$_6$;HexNAc$_2$Hex$_7$;HexNAc$_2$Hex$_8$;HexNAc$_2$Hex$_9$ |
| **N607AS** | α | **Yes** | 97.87% | HexNAc$_2$Hex$_5$;HexNAc$_2$Hex$_6$;HexNAc$_2$Hex$_8$;HexNAc$_2$Hex$_9$;HexNAc$_3$Hex$_5$**NeuAc**;HexNAc$_3$Hex$_6$;HexNAc$_4$Hex$_5$NeuAc;HexNAc$_4$Hex$_5$Fuc;HexNAc$_5$Hex$_5$;HexNAc$_5$Hex$_5$Fuc |
| N764IT | β | No | 53.85% | HexNAc$_2$Hex$_8$;HexNAc$_2$Hex$_9$ |
| N900YT | β | | 100% | HexNAc$_2$Hex$_5$;HexNAc$_2$Hex$_6$;HexNAc$_2$Hex$_7$;HexNAc$_2$Hex$_8$;HexNAc$_2$Hex$_9$;HexNAc$_3$Hex$_5$Fuc;HexNAc$_3$Hex$_6$ |
| **N913GS** | β | **Yes** | 100% | HexNAc$_3$Hex$_5$**NeuAc**;HexNAc$_4$Hex$_5$**NeuAc** |

glycosylation frequency=number of glycosylated peptide/number of total peptide; HexNAc, N-acetylhexosamine; Hex, hexose; Fuc, fucose; **NeuAc, N-acetylneuraminic acid (sialic acid)**.

**Table S2. List of site-specific N-glycans of IGF1R identified from glycoproteomic analysis**
